# Optimizing interneuron circuits for compartment-specific feedback inhibition

**DOI:** 10.1101/2020.11.17.386920

**Authors:** Joram Keijser, Henning Sprekeler

## Abstract

Cortical circuits process information by rich recurrent interactions between excitatory neurons and inhibitory interneurons. One of the prime functions of interneurons is to stabilize the circuit by feedback inhibition, but the level of specificity on which inhibitory feedback operates is not fully resolved. We hypothesized that inhibitory circuits could enable separate feedback control loops for different synaptic input streams, by means of specific feedback inhibition to different neuronal compartments. To investigate this hypothesis, we adopted an optimization approach. Leveraging recent advances in training spiking network models, we optimized the connectivity and short-term plasticity of interneuron circuits for compartment-specific feedback inhibition onto pyramidal neurons. Over the course of the optimization, the interneurons diversified into two classes that resembled parvalbumin (PV) and somatostatin (SST) expressing interneurons. The resulting circuit can be understood as a neural decoder that inverts the nonlinear biophysical computations performed within the pyramidal cells. Our model provides a proof of concept for studying structure-function relations in cortical circuits by a combination of gradient-based optimization and biologically plausible phenomenological models.

## Introduction

Cortical inhibitory interneurons vary dramatically in shape, gene expression pattern, electrophysiological and synaptic properties and in their downstream targets [1]. Some cell types, e.g., somatostatin (SST)-positive interneurons [2] and some neurogliaform cells in layer 1 [3], predominantly project to pyramidal cell (PC) dendrites. Others—e.g., parvalbumin positive (PV) basket and chandelier cells—primarily inhibit the peri-somatic domain of PCs [4]. Some interneurons receive depressing synapses from PCs, others facilitating synapses [5, 6]. But what is the function of these differences?

One of inhibition’s core functions is to prevent run-away excitation [7] by means of feedback inhibition that tracks excitatory inputs. This has led to the concept of excitation-inhibition (E/I) balance [8], i.e., the idea that strong excitatory currents are compensated by inhibitory currents of comparable size. E/I balance is thought to shape cortical dynamics [8, 9] and computations [10, 11] and can be established by means of inhibitory forms of plasticity [12, 13, 14]. Selective disruptions of E/I balance are thought to play a key role during learning [15], while chronic disturbances have been implicated with psychiatric diseases, including autism [16, 17] and schizophrenia [18, 19].

Originally conceived as a balance on average [8], E/I balance turned out to be specific to sensory stimuli [20, 21], in time [22, 23], across neurons [24] and to neural activation patterns [25]. The number of excitatory and inhibitory synapses could even be balanced at the subcellular level[26], in a cell-type specific way [27]). Given this high specificity, we hypothesized that excitation and inhibition also balance individually in different neuronal compartments, and that this could be mediated at least in part by compartment-specific feedback inhibition.

Different neuronal compartments often receive input from different sources [28] and integrate these inputs nonlinearly by means of complex cellular dynamics [29, 30]. For example, the apical dendrites of L5 pyramidal cells (PCs) can generate nonlinear calcium events in response to coincident somatic and dendritic inputs [31]. Hence, neuronal output spike trains can have a complex nonlinear dependence on the inputs arriving in different compartments. This poses a challenge for compartment-specific feedback inhibition, which would require interneurons to invert the nonlinear dependence by recovering local dendritic input from pyramidal output. It is therefore far from clear that a compartment-specific feedback inhibition can be achieved at all by means of biologically plausible circuits. If it can, however, it would have to rely on an interneuron circuit that is closely matched to the electrophysiological properties of the cells it inhibits. Parts of the complexity of cortical interneuron circuits could then be interpreted in light of the intrinsic properties of PCs.

Unfortunately, the nature of such a correspondence between the electrophysiology of inhibited cells and suitable interneuron circuits is far from obvious. We reasoned that we could gain insights by means of a model-based optimization approach, in which interneuron circuits are optimized for feedback inhibition onto pyramidal cells with given biophysical properties. Here, we illustrate this ansatz by optimizing interneuron circuits for a nonlinear two-compartment model of L5 pyramidal cells [32]. We show that over the course of the optimization, an initially homogeneous interneuron population diversifies into two classes, which share many features of cortical PV and SST interneurons. One class primarily inhibits the somatic compartment of the PCs and receives depressing synaptic inputs. The other class primarily inhibits PC dendrites and received facilitating inputs. We show how this diversification can be understood from an encoding-decoding perspective, in which the biophysics of the PCs encode two different input streams in a multiplexed code [33], which is in turn decoded by the interneuron circuit. These findings support the idea that parts of the complexity of cortical interneuron circuits could be interpreted in light of the intrinsic properties of PCs and illustrate how modeling could provide a means of unravelling these interdependencies between the cellular and the circuit level.

## Results

To investigate which aspects of cortical interneuron circuits can be understood from the perspective of compartment-specific inhibition, we studied a spiking network model comprising pyramidal cells (PCs) and interneurons (INs) (see Methods). PCs were described by a two-compartment model consisting of a soma and an apical dendrite. The parameters of this model were previously fitted to capture dendrite-dependent bursting [32]. PCs received time-varying inputs in both the somatic and the dendritic compartment, and inhibitory inputs from INs. INs were described by an integrate-and-fire model. They received excitatory inputs from the PCs, and inhibitory inputs from other INs.

We optimized the interneuron circuit for a compartment-specific feedback inhibition. In the presence of time-varying external input, feedback inhibition tracks excitatory inputs in time [8, 23]. We therefore enforced compartment-specific feedback inhibition by minimizing the mean squared error between excitatory and inhibitory inputs in both compartments, by means of gradient descent with surrogate gradients [34]. Importantly, we optimized not only the strength of all synaptic connections in the network, but also the short-term plasticity of the PC → IN connections (see Methods).

### Interneuron diversity emerges during optimization

Before the optimization, interneurons formed a single, homogeneous group (Fig. 1a, top). Most inhibited both somatic and dendritic compartments (Fig. 1b, top) and PC → IN connections showed non-specific synaptic dynamics (Fig. 1c, top). Moreover, excitation and inhibition were poorly correlated, particularly in the dendrite (Pearson correlation coefficients 0.49 (soma) & 0.08 (dendrite)), suggesting that the network did not generate compartment-specific feedback inhibition (Fig. 1d, top).

**Figure 1:**
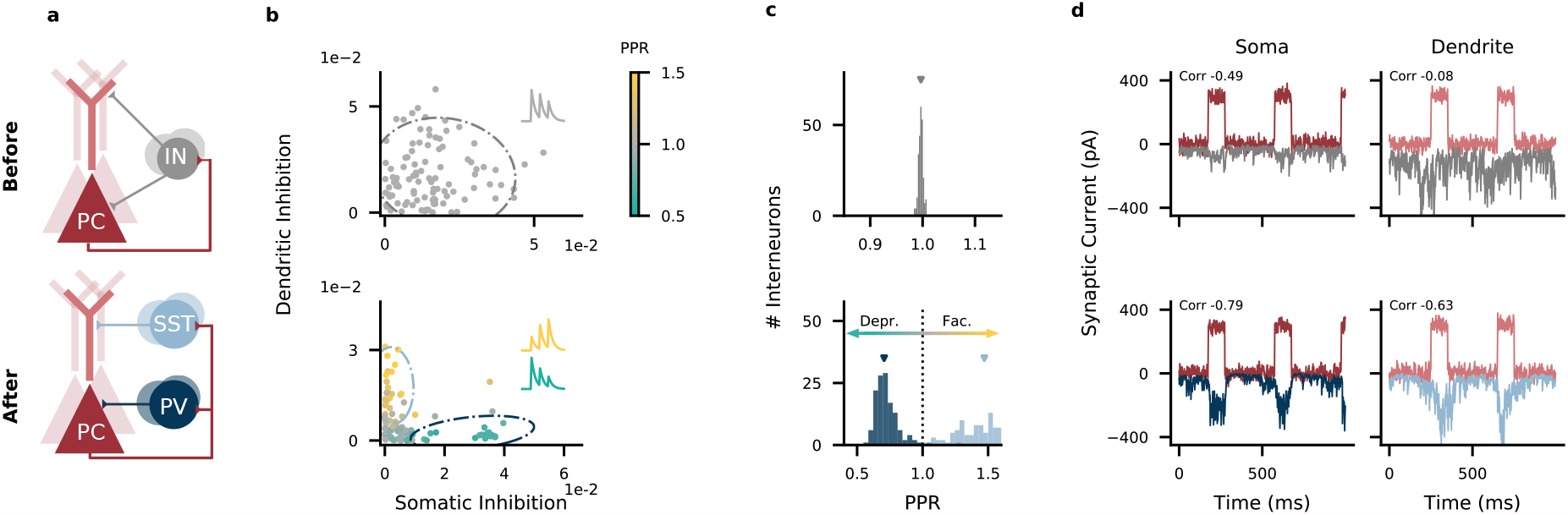
Interneuron diversity emerges in networks optimized for compartment-specific inhibition. **(a)** Network structure before (top) and after optimization (bottom). PC, pyramidal cell; IN, interneuron; PV, parvalbumin-positive IN; SST, somatostatin-positive IN. Recurrent inhibitory connections among INs omitted for clarity. **(b)** Strength of somatic and dendritic inhibition from individual INs. Dashed lines: 95% density of a Gaussian distribution (top) and mixture of two Gaussian distributions (bottom) fitted to the connectivity and Paired Pulse Ratio (PPR) data of 5 networks (marginalized over PPR). **(c)** PPR distribution (data from 5 networks). Mean PPR before optimization: 1.00; after optimization: 0.73 (PV cluster, *n* = 133) and 1.45 (SST cluster, *n* = 113). **(d)** Excitatory (red) and inhibitory (top: gray, bottom: blue) currents onto PC compartments (average across *N*_*E*_ = 400 PCs). Inset: correlation between compartment-specific excitation and inhibition.

During optimization, the interneurons split into two groups (Fig. 1a, bottom) with distinct connectivity (Fig. 1b, bottom; see also Connectivity among interneurons) and short-term plasticity (Fig. 1c, bottom). One group received short-term depressing inputs from PCs and preferentially targeted their somatic compartment, akin to PV interneurons. The other group received short-term facilitating inputs from PCs and targeted their dendritic compartment, akin to SST interneurons. For simplicity, we will henceforth denote the two interneuron groups as PV and SST interneurons. After the optimization, excitation and inhibition were positively correlated in both compartments (Pearson correlation coefficients 0.79 (soma) & 0.63 (dendrite); Fig. 1d, bottom). Note that the E/I balance is slightly less tight in time in the dendrites than in the somata (Fig. 1d), because synaptic short-term facilitation causes a delay in the signal transmission between PCs and SST interneurons [35, more details below].

To confirm the benefit of two non-overlapping interneuron classes, we performed control simulations in which each interneuron was pre-assigned to target either the soma or the dendrite, while synaptic strengths and short-term plasticity were optimized. Consistent with a benefit of a specialization, the correlation of excitation and inhibition in the two compartments was as high as in fully self-organized networks (Fig. 2). Optimized networks with pre-assigned interneuron classes also showed the same diversification in their short-term plasticity, resembling that of PV and SST neurons (Figs. 2, A.1).

**Figure 2:**
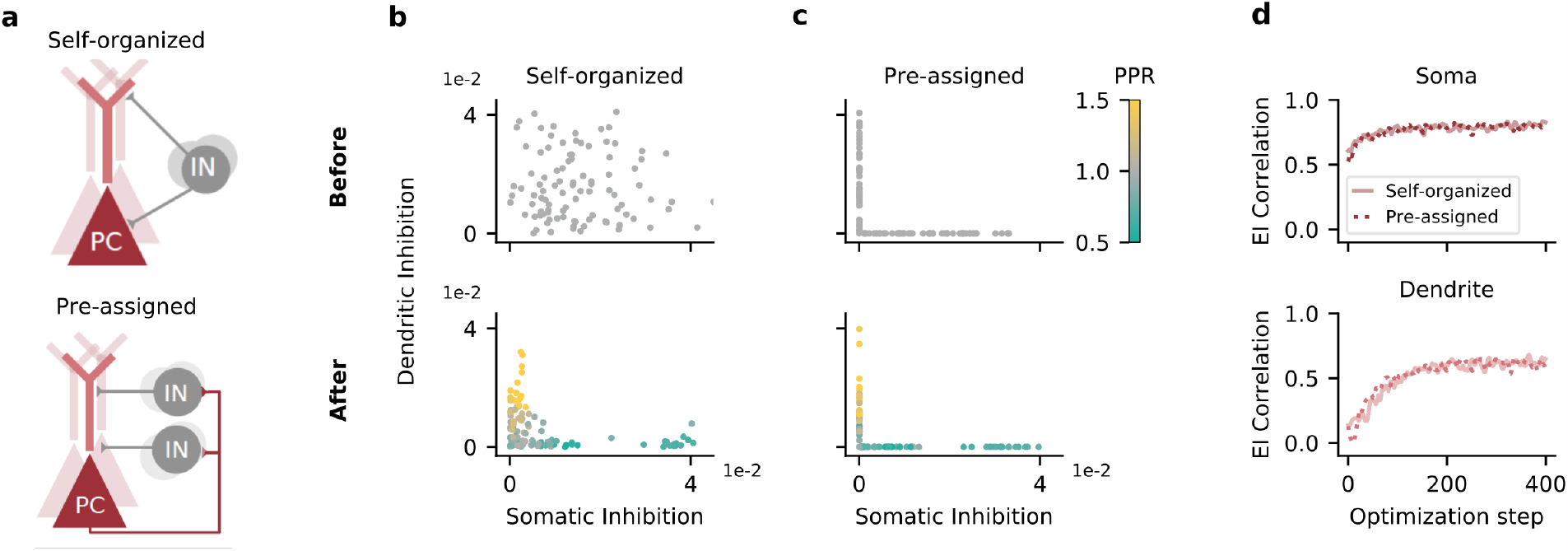
Compartment-assigned interneurons develop into PV- and SST-like populations. **(a)** Circuit before learning. Top, interneurons (INs) can inhibit both compartments of principal cells (PCs) and need to self-organize, as in Fig. 1. Bottom, INs are pre-assigned to inhibit a single PC compartment. **(b)** IN→PC weights before (top) and after (bottom) optimization. Interneurons self-organize into a population that preferentially inhibits the soma, and a population that preferentially inhibits the dendrites. Data differs from Fig. 1 due to random parameter initialization and sampling of training data. **(c)** As (b), but with interneurons randomly assigned to inhibit a single compartment (soma or dendrite). Mean PPR: 0.72 (soma-inhibiting population), 1.17 (dendrite-inhibiting population). **(d)** Correlation between compartment-specific excitation and inhibition over the course of the optimization. Solid line: INs were not assigned to a single compartment (Self-organized). Dashed line: INs were assigned to a single compartment (Pre-assigned). Data is smoothed with a Gaussian kernel (width: 2).

### Feedback inhibition decodes compartment-specific inputs

For compartment-specific feedback inhibition, the interneuron circuit has to retrieve the somatic and dendritic input to PCs from the spiking activity of the PCs. This amounts to inverting the nonlinear integration performed in the PCs (Fig. 3a). How does the circuit achieve this? Recently, it was proposed that the electrophysiological properties of PCs support a multiplexed neural code that simultaneously represents somatic and dendritic inputs in temporal spike patterns ([33], Fig. 3b). In this code, somatic input increases the number of events, where events can either be single spikes or bursts (see Methods). Dendritic input in turn increases the probability that a somatic spike is converted into a burst (burst probability). Providing soma- or dendrite-specific inhibition then amounts to decoding the event rate or burst probability, respectively. Such a decoding can be achieved in circuits with short-term plasticity and feedforward inhibition [33], and we expected that our network arrived at a similar decoding scheme.

**Figure 3:**
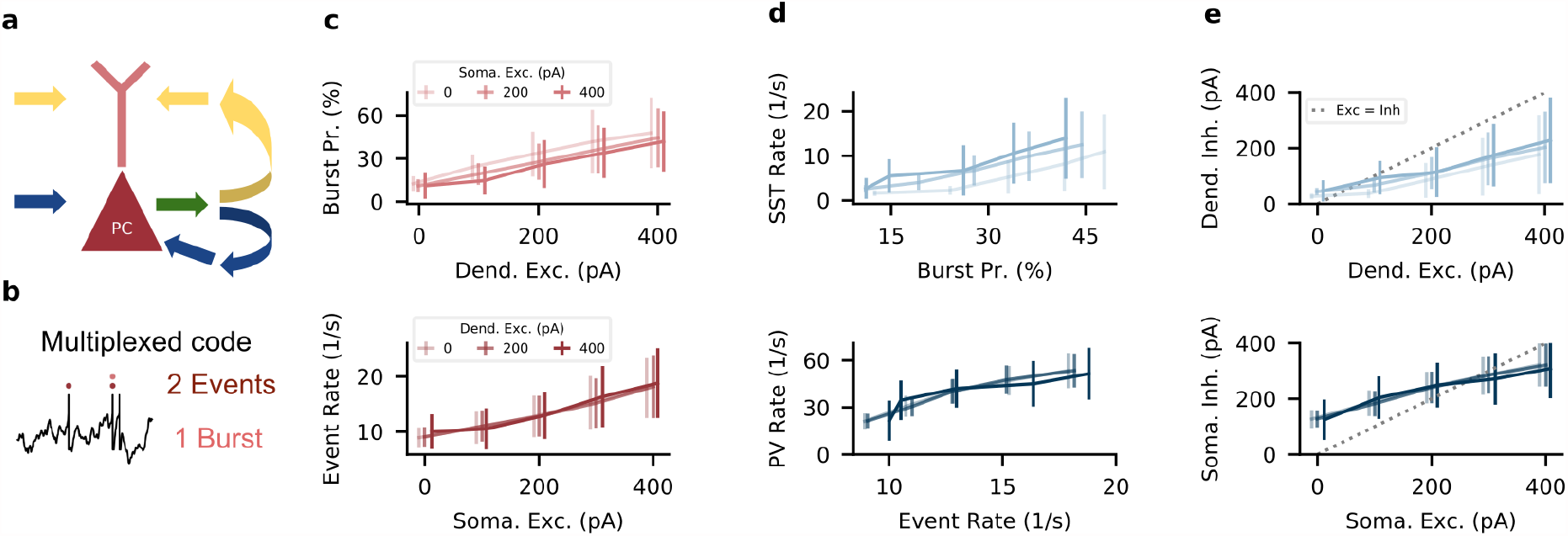
The interneuron circuit decodes somatic and dendritic inputs to PCs. **(a)** PC somata and dendrites receive uncorrelated input streams (yellow and blue) that, from PC output spikes (green), have to be separated into compartment-specific inhibition (yellow and blue). **(b)** PCs use a multiplexed neural code. Somatic input leads to events (singlets or bursts). Dendritic input converts singlets into bursts. **(c)** Top: Excitatory input to PC dendrites increases burst probability. In this and other top panels (d,e), the shading indicates strength of background somatic input, and error bars indicate sd over 10 stimulus repetitions. Bottom: Excitatory input to PC somata increases event rate. In this and other bottom panels (d,e), the shading indicates strength of background dendritic input. **(d)** Top: SST rate increases with bursts probability. Bottom: PV rate increases with PC events. **(e)** Top: dendritic inhibition increases with dendritic excitation, but is only weakly modulated by somatic excitation. Positions on *x*-axis are shifted by 10 pA for visual clarity, error bars indicate sd during 10 stimulus repetitions. Bottom: somatic inhibition increases with somatic excitation, but is invariant to dendritic excitation.

We tested this hypothesis by injecting current pulses to PC somata and dendrites (see Methods). Stronger dendritic input increased the burst probability, which increased the firing rate of SST interneurons via facilitating synapses. The increased SST rate increased dendritic inhibition (Fig. 3c-e, top). Analogously, stronger somatic input increased the event rate, which increased the firing rate of PV interneurons via depressing synapses. The increased PV rate increased somatic inhibition (Fig. 3c-e, bottom). Importantly, inhibition was specific to each compartment (shaded lines indicate input strength to the other compartment): Because PV interneurons were selectively activated by PC events, somatic inhibition was largely unaffected by dendritic excitation. Similarly, SST interneurons were selectively activated by PC bursts, such that dendritic inhibition was largely unaffected by somatic excitation. In the model, interneurons therefore provide compartment-specific inhibition by demultiplexing the neural code used by the PCs.

### Effect of correlations between somatic and dendritic input

So far we assumed that PC somata and dendrites receive uncorrelated input. Recent work, however, suggests that somatic and dendritic activity are correlated [36, 37], potentially reducing the need for compartment-specific inhibition. We therefore tested how correlated inputs affect interneuron specialization by optimizing separate networks for different input correlations. We found that increasing correlation between somatic and dendritic inputs gradually reduced the separation between the interneuron classes (Fig. 4a,b). For high input correlation, optimized networks contained a continuum in their connectivity and short-term plasticity (Fig. 4a,b). However, the presence of short-term plasticity was necessary for a dendritic E/I balance for a range of input correlations (Fig. 4c). At high correlations, somatic and dendritic inputs are sufficiently similar to make the effect of short-term facilitation negligible. Note that although in this case, distinct interneuron populations were not necessary, the presence of IN classes was also not harmful for E/I balance. A pre-assignment of the interneurons into classes maintained the E/I correlation in both compartments and for any correlation level (Fig. A.1). Finally, we found that interneuron specialization degraded with increasing baseline activity of the INs (Fig. A.2), because high firing rates allow non-specialized inhibition to cancel out (see mathematical analysis in Supplementary Materials. However, a pre-assignment of interneurons into classes again maintained the E/I correlation for different baseline activity levels (Fig. A.1).

**Figure 4:**
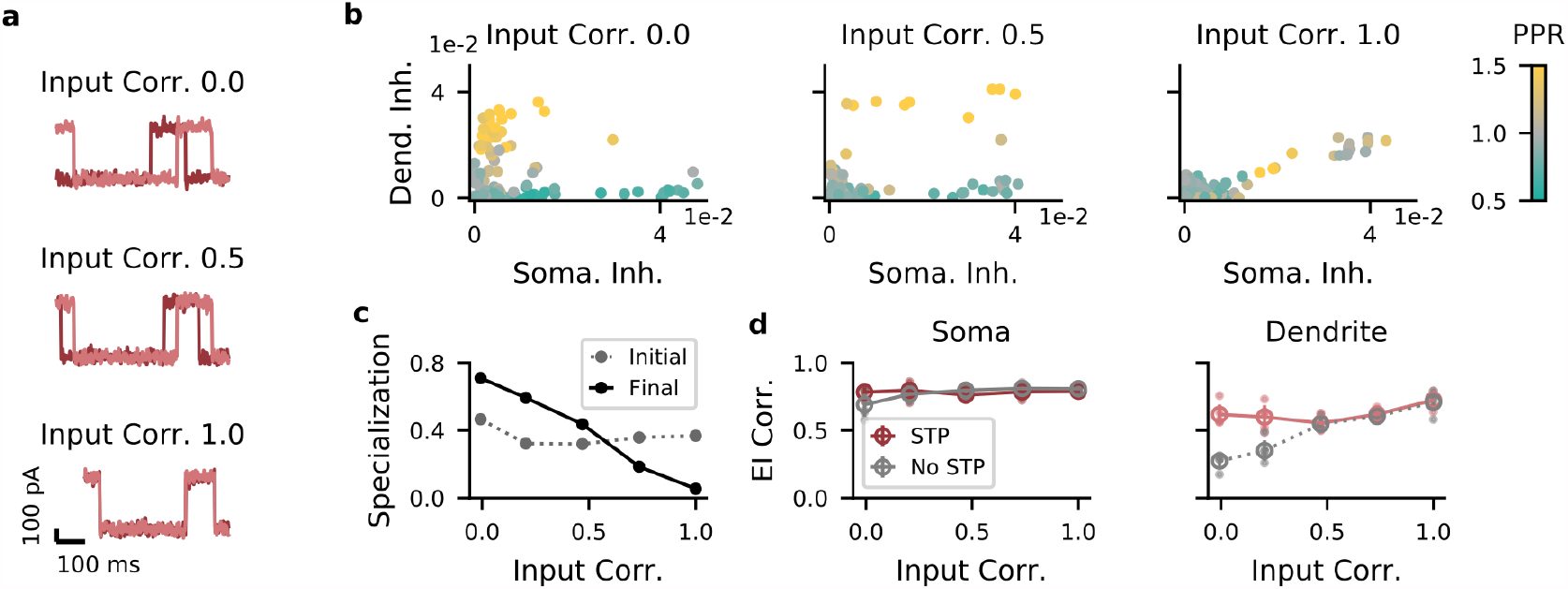
Correlations between dendritic and somatic input reduce interneuron specialization. **(a)** Examples for synaptic traces corresponding to different correlation levels. Dark red, somatic current; light red, dendritic input. **(b)** Strength of somatic vs. dendritic inhibition from all INs. Left, middle, right: input correlation coefficient 0 (low), 0.5 (medium), and 1 (high), respectively. **(c)** Specialization of IN → E weights. If each IN targets either soma or dendrites, the specialization is 1 (see Methods). Gray: specialization of initial random network; black: specialization after optimization. **(d)** Left: In the soma, excitation and inhibition are balanced across a broad range of input correlations, with or without short-term plasticity (STP). Right: In the dendrites, excitation and inhibition are balanced only with STP when input correlations are small.

### Connectivity among interneurons

Because interneurons subtypes also differ in their connectivity to other interneurons [38, 39], we included IN → IN synapses in our optimization. After classifying INs as putative PV and SST neurons using a binary Gaussian mixture model, we found that the connections between the interneuron classes varied systematically in strength. While PV ↔ PV connections, PV → SST connections and SST ↔ SST connections were similar in strength on average, SST → PV were consistently stronger (Fig. 5a), presumably to compensate for the relatively low SST rates (Fig. 3d).

**Figure 5:**
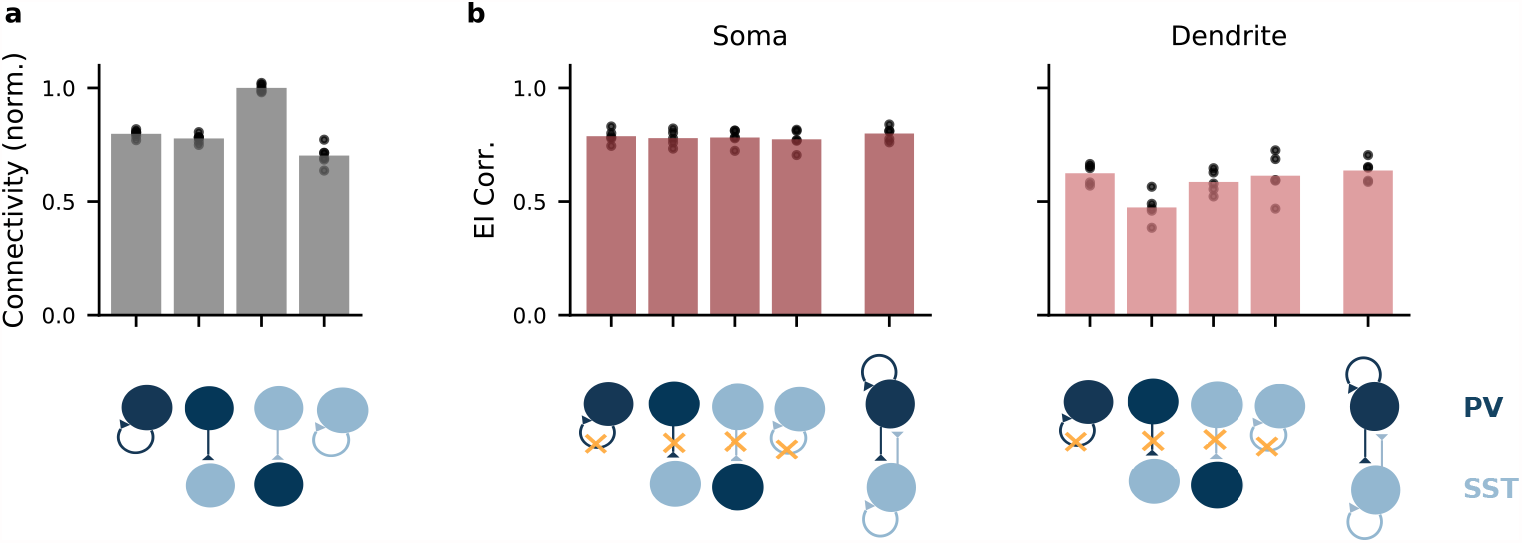
Recurrent inhibitory connectivity after learning. **(a)** Connectivity between IN populations. From left to right: PV↔PV, PV→SST, SST→PV, SST↔SST. Bars indicate mean over all networks, dots indicate individual networks. **(b)** Performance as measured by the correlation between excitation and inhibition to PC soma (left) and dendrites (right) of networks optimized lacking specific connections. Data at the very right: EI correlation in network with unconstrained connectivity. Only loss of PV → SST connectivity has a clear effect on dendritic EI correlations. Open circles, mean over 5 batches of 8 stimuli with random amplitudes. Small filled circles, individual batches.

To investigate which connections were necessary, we simulated knockout experiments in networks with pre-assigned interneuron classes, in which we removed individual connections types. We found that only PV → SST connections were necessary for a dendritic E/I balance (Fig. 5b). Note that although earlier work did not find PV → SST connectivity in the primary visual cortex of young mice [38], these connections seem to be present in primary visual and somatosensory cortex of older animals [39, 40].

To understand the role of the different IN→IN connections, we performed a mathematical analysis of a simplified network model. The model also contains a population of principal cells (PC) and two populations of interneurons corresponding to PV and SST interneurons, but in contrast to the spiking model, neural activities are represented by continuously-varying rates. The population rates of PV and SST interneurons are denoted by *p* and *s*, respectively. The activity of PCs is described by two rates: an event rate *e* that is driven by somatic input and a burst rate *b* that is driven by dendritic input. The short-term plasticity of a given synapse type is characterized by a single, static parameter, which characterizes the relative efficiency at which events and bursts are transmitted. Synapses for which this parameter is 1 transmit events but not bursts, i.e., they are “perfectly depressing”. Synapses for which this parameter is 0 transmit only bursts, i.e., they are “perfectly facilitating”. These assumptions allowed us to mathematically analyze the interneuron connectivity required for compartment-specific feedback inhibition. We will only summarize the results, the full analysis is described in Supplementary Materials.

Let us first consider the case of dendritic feedback inhibition. The model states that the activity *s* of the SST neurons is given by a linear combination of the event and burst rate: *s* = *Ae* + *B b*, with factors *A, B* that depend on the connectivity and short-term plasticity in the circuit in a complicated way. If we assume that SST interneurons target exclusively the PC dendrites, compartment-specific feedback inhibition requires that the activity of SST interneurons depends on dendritic but not somatic input to PCs. Because those two inputs drive the event rate and burst rate, respectively, this condition reduces to the mathematical condition that *A* = 0. Using the dependence of *A* on the circuit parameters (see Supplementary Materials), we get the condition

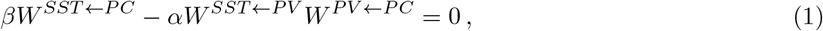

where *W* ^*Y ←X*^ denotes the strength of the synaptic connection between population *X* and *Y*. The two parameters *α, β* are the short-term plasticity parameters and quantify how well events are transmitted via the PC→PV and PC→SST connections, respectively.

Condition [1] has an intuitive interpretation. The first term describes how much somatic PC input influences SST activity via the monosynaptic pathway PC → SST. The second term corresponds to the disynaptic pathway PC → PV → SST. The condition therefore states that unless PC→ SST connections are “perfectly facilitating” (*β* = 0), the disynaptic PC → PV → SST pathway is necessary (Fig. 5) to avoid that somatic input generates dendritic inhibition. The observation that a knock-out of these connections reduces the dendritic E/I correlation in the spiking network (Fig. 5b) can therefore be understood as a result of an imperfect facilitation in the PC→SST connection. Indeed, we observed that the Tsodyks-Markram model [41] we used to describe the short-term plasticity in the spiking network cannot achieve a perfectly facilitating synapse in the presence of ongoing activity, even for an initial release probability *U* = 0, because preceding spikes always leave behind a residual level of synaptic facilitation.

An analogous analysis suggests that disynaptic PC → SST→ PV inhibition is necessary to prevent dendritic inputs from generating somatic inhibition (Supplementary Materials), providing a possible function of experimentally observed SST → PV connectivity. At first sight, this appears in conflict with the observation that a knock-out of this connection did not reduce the E/I balance in the soma. However, because bursts are comparatively rare [33], event rate and overall firing (including additional spikes in bursts) are highly correlated. Therefore, the overall firing rate is a good proxy for somatic input and imperfections in synaptic depression in the PC→PV connection do not introduce a sufficiently large problem to necessitate feedforward inhibition via the PC→SST→PV pathway.

## Discussion

Feedback inhibition ensures the stability of cortical circuits [42, 43, 11, 44]. Our model indicates that this feedback could operate on a level as fine-grained as different cellular compartments receiving different input streams, and that the required circuitry bears similarity to the one observed in cortex. In particular, we found that an optimization for feedback inhibition led to the emergence of two inhibitory cell classes that resemble PV and SST interneurons in their connectivity and short-term plasticity. This diversification was robust to correlations between somatic and dendritic input, although increasing correlations prompted the SST-like model neurons to contact not only the dendritic, but also the somatic compartment. This is consistent with the extensive branching of cortical SST neurons within the layer that contains their cell body [2]. Even in cases in which the gradient-based optimization did not drive a clear division into cell classes, an artificial pre-assignment of the interneurons did not impair the feedback inhibition.

We would like to emphasize that while we optimized for feedback inhibition in different neuronal compartments, the model operates on an ensemble level in the sense that all neurons in the network received the two same time-varying signals in their soma and dendrite. This allows the interneurons to use event or burst rates of the whole ensemble to infer somatic and dendritic inputs with high temporal fidelity [33]. The question of the specificity of feedback inhibition on the population level is an orthogonal one and not fully resolved. The dense and seemingly unspecific connectivity of many interneurons [45, 46] suggests that feedback inhibition operates on the level of the local population, blissfully ignoring the functional identity of the neurons it targets [47]. More recent results have indicated a correlation between the sensory tuning and the synaptic efficacy of interneuron-pyramidal cell connections, however, suggesting that feedback inhibition could operate on the level of functionally identified ensembles [48, 13]. A natural extension of this work would be to endow the pyramidal cells with a tuning to different somatic and dendritic input streams and thereby define functional ensembles. Notably, the ensemble affiliation of a given neuron may differ for soma and dendrite, e.g., two populations of neurons could receive distinct somatic, but identical dendritic inputs. How this would be reflected in the associated feedback-optimized interneuron circuit is an interesting question, but beyond the scope of the present work.

A natural question for optimization-based approaches is how the optimization can be performed by biologically plausible mechanisms. The gradient-based optimization we performed relies on surrogate gradients [34, 49] and a highly non-local backpropagation of errors both through the network and through time [50, 51], mechanisms that are unlikely implemented verbatim in the circuit [52]. We think of the suggested optimization approach rather as a means to understanding functional relations between different features of neural circuits, i.e., the relation between the biophysics of pyramidal cells and the surrounding interneuron circuits. At this point, we prefer to remain agnostic as to the mechanisms that establish these relations. While an activity-dependent refinement of the circuit is likely, the diversification of the interneurons into PV and SST neurons is clearly not driven by activity-dependent mechanisms alone [53]. SST Martinotti cells migrate to the embryonic cortex via the marginal zone, while PV basket cells migrate via the subventricular zone [54]. Their identity is hence determined long before they are integrated into functional circuits. These developmental programs are likely old on evolutionary time scales given that interneuron diversity seems highly conserved [55, 56]. Yet, even when interpreted as an evolutionary optimization, interneuron circuits probably did not evolve to perform feedback inhibition for pre-existing biophysical properties of pyramidal neurons. Such systematic relations between different circuit features are more likely a result of co-evolution, where the different features mutually enabled the successful selection of the others.

Given these considerations, we refrain from predictions regarding the optimization process. Still, the model can make predictions regarding the nature of the optimized state. First, it predicts that PV and SST rates correlate primarily with somatic and dendritic activity, respectively. Second, inhibiting SST neurons should increase PC bursting, as observed in hippocampus [57] and cortex [58]. The role of short-term facilitation could be tested by silencing the necessary gene Elfn1 [59, 60]. On a higher level, the model suggests a relation between the biophysical properties of excitatory neurons and the surrounding interneuron circuit. This is consistent, e.g., with the finding that the prevalence of pyramidal cells and dendrite-targeting Martinotti cells seems to be correlated across brain regions [61].

While the synaptic targets and the incoming short-term plasticity of the two emerging interneuron classes are similar to those of PV and SST interneurons, the optimized inhibitory circuitry is not a perfect image of cortex. Aside from the obvious incompleteness in terms of other interneuron types, other features, such as the often observed weak connectivity from PV to SST neurons [38] did not result from the optimization (Fig. 5). However, even if our assumption that the interneuron circuit performs compartment-specific feedback inhibition was correct, a perfect match to cortex is probably not to be expected. Firstly, the pyramidal cell model we used is clearly a very reduced depiction of a real pyramidal cell. Because the inhibitory circuitry is optimized for the nonlinear processing performed by these cells, anything that is wrong in the pyramidal cell model will also be wrong in the optimized circuit. It will be interesting to see how the suggested optimization framework generalizes to computations performed by more complex neuronal morphologies [30]. Secondly, the optimized circuitry is also sensitive to other modelling choices. For example, the circuit separates spikes and bursts by a synergy between short-term plasticity and interneuron connectivity. A wrong short-term plasticity model will therefore lead to a wrong connectivity in the circuit. Here, it will be interesting to see how a more expressive model of short-term plasticity [62] influences the optimal circuit structure. Finally, of course, our optimality assumption could be wrong to different degrees. We could be wrong in detail: Even if the idea of compartment-specific feedback inhibition was correct, our mathematical representation thereof – matching excitation and inhibition in time – could be wrong, with corresponding repercussions in the optimized circuit. Or we could be wrong altogether: PV and SST interneurons serve an altogether different function, and feedback inhibition is merely a means to a completely different end, such as behavioral circuit modulation [58, 63] or the control of plasticity [15, 64].

Notwithstanding the dependence of the final circuit on specific model choices, we believe that the suggested optimization approach provides a broadly applicable schema for analyses of structure-function relations of interneuron circuits. On a coarser level of biological detail, optimization approaches have recently been quite successful at linking abstract computations to the neural network level [65, 66, 67]. While similar in spirit, our approach takes this optimization ansatz from the level of dynamical systems analyses of rate-based recurrent neural networks to the detailed level of spiking circuits with multi-compartment neurons and short-term plasticity. It will be exciting to see how biological mechanisms on this level of detail support more advanced computations than the mere stabilization of the circuit considered here, but that is clearly a larger research program that extends well beyond the proof of concept presented here.

## Acknowledgements

J.K. was supported by a PhD scholarship from the Einstein Center for Neurosciences Berlin. We thank Loreen Hertäg, Basile Confavreux, Laura Bella Naumann, Filip Vercruysse, and Robert Tjarko Lange for comments on earlier drafts.

## Methods

### Network Model

We simulated a spiking network model consisting of *N*_*E*_ pyramidal cells (PCs) and *N*_*I*_ interneurons (INs), as in earlier work [33]. PCs are described by a two-compartment model [32]. The membrane potential *v*^*s*^ in the somatic compartment is modeled as a leaky integrate-and-fire unit with spike-triggered adaptation:

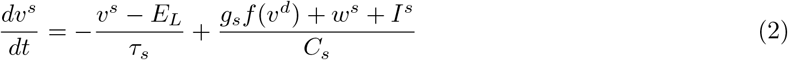

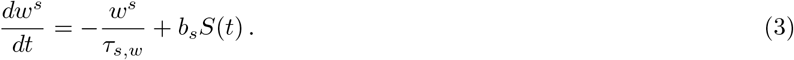

Here, *E*_*L*_ denotes the resting potential, *τ*_*s*_ the membrane time constant and *C*_*s*_ the capacitance of the soma. *I*^*s*^ is the external input, and *w*^*s*^ the adaptation variable, which follows leaky dynamics with time constant *τ*_*s,w*_, driven by the spike train *S* emitted by the soma. *b*_*s*_ controls the strength of the spike-triggered adaptation. *v*^*d*^ is the dendritic membrane potential, the conductance *g*_*s*_ controls how strongly the dendrite drives the soma, and *f* the nonlinear activation of the dendrite:

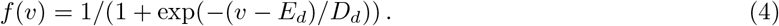

The half-point *E*_*d*_ and slope *D* of the transfer function *f* control the excitability of the dendrite. When the membrane potential reaches the spiking threshold *ϑ*, it is reset to the resting potential and the PC emits a spike. Every spike is followed by an absolute refractory period of *τ*_*r*_.

The dynamics of the dendritic compartment are given by:

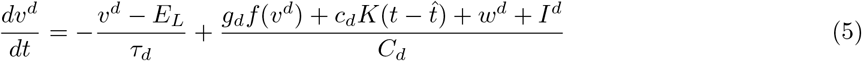

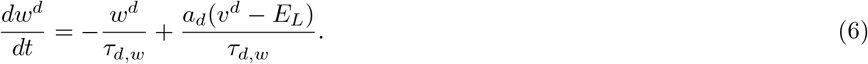

In addition to leaky membrane potential dynamics with time constant *τ*_*d*_, the dendrite shows a voltage-dependent nonlinear activation *f*, the strength of which is controlled by *g*_*d*_. This nonlinearity allows the generation of dendritic plateau potentials (“calcium spikes”). Somatic spikes trigger backpropagating action potentials in the dendrite, modeled in the form of a boxcar kernel *K*, which starts 1ms after the spike and lasts 2ms. The amplitude of the backpropagating action potential is controlled by the parameter *c*_*d*_. The dendrite is subject to a voltage-activated adaptation current *w*^*d*^, which limits the duration of the plateau potential. This adaptation follows leaky dynamics with time constant *τ*_*d,w*_. The strength of the adaptation is given by the parameter *a*_*d*_. Note that the model excludes sub-threshold coupling from the soma to the dendrite.

The interneurons are modeled as leaky integrate-and-fire neurons:

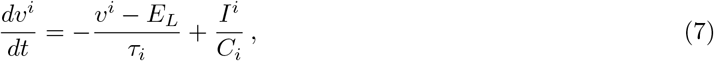

with time constant *τ*_*i*_. Spike threshold, resting and reset potential, and refractory period are the same as for the PCs.

All neurons receive an external background current to ensure uncorrelated activity, which follows Ornstein-Uhlenbeck dynamics

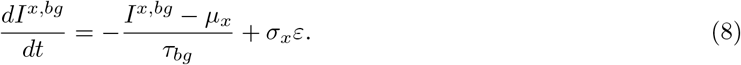

Here, *x* ∈ {*s, d, i*} refers to the soma, dendrite, or interneuron, respectively, and *ε* is standard Gaussian white noise with zero mean and correlation ⟨ *ε*(*t*)*ε*(*t*′) ⟩ = *δ*(*t* − *t*′).

In addition, the somatic and dendritic compartments received step currents mimicking external signals (see Optimization), as well as recurrent inhibitory inputs. The recurrent input to compartment *x* ∈ {*s, d*} of the *i*th principal cell was given by

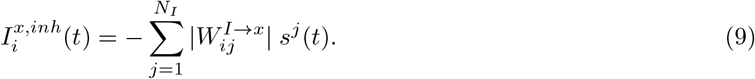

where *s*^*j*^ is the synaptic trace that is increased at each presynaptic spike and decays with time constant *τ*_*syn*_ otherwise:

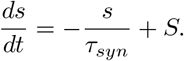

The compartment-specific inhibitory weight matrices *W* ^*I→x*^, *x* ∈ {*s, d*} were optimized; the absolute value in Eq. 9 ensured positive weights.

The recurrent input to the *i*th interneuron was given by:

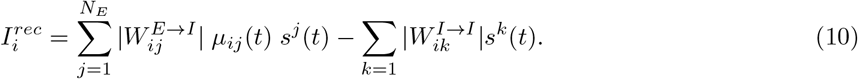

The function *µ*_*ij*_(*t*) implements short-term plasticity according to the Tsodyks-Markram model [41]. *µ*(*t*) is the product of a utilization variable *u* and a recovery variable *R* that obey the dynamics

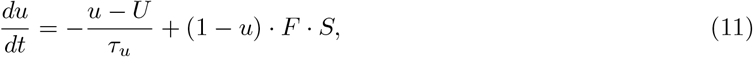

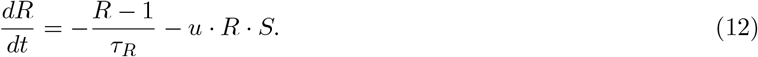

*U* is the initial release probability, which is optimized by gradient descent. *F* is the facilitation fraction, and *τ*_*R*_, *τ*_*u*_ are the time constants of facilitation and depression, respectively. All parameter values are listed in Table 1 (Supplementary Materials)

**Table 1:** Parameter values related to network simulation

| Symbol | Value | Unit | Description |
| --- | --- | --- | --- |
| $N_E$ | 400 | - | Number of exc. neurons |
| $N_I$ | 100 | - | Number of inh. neurons |
| $E_L$ | -70 | mV | reversal and reset potential |
| $\vartheta$ | -50 | mV | spiking threshold |
| $\tau_{s/d/i}$ | 16 / 7 / 10 | ms | time const. soma/ dend./inh. membrane |
| $\tau_r$ | 3 | ms | refractory time soma and inh. |
| $g_{s/d}$ | 1300 / 1200 | pA | Coupling from dend to soma |
| $C_{s/d/i}$ | 370 / 170 / 100 | pF | Conductance of soma/dend./inh. |
| $\tau_{s/d,w}$ | 100 / 30 | ms | Time const. adaptation soma/dend. |
| $b_s$ | -200 | pA | Spike-triggered adaptation (soma) |
| $a_d$ | -13 | nS | Voltage-driven adaptation (dend) |
| $c_d$ | 2600 | pA | Coupling soma to dend. |
| $E_d$ | -38 | mV | position dend. nonlinearity |
| $D_d$ | 6 | mV | steepness of dend. nonlinearity |
| $\mu_{s/d/i}$ | 400 / -300 / -100 | pA | mean background input soma/dend./inh. |
| $\sigma_{s/d/i}$ | 450 / 450 / 400 | pA | sd background input |
| $\tau_{bg}$ | 2 | ms | time const. background input |
| $\tau_{syn}$ | 5 | ms | time const. synapses |
| $\tau_u$ | 100 | ms | time const. facilitation |
| $\tau_R$ | 100 | ms | time const. depression |
| $F$ | 0.1 | - | facilitation jump |

Finally, the network parameters were scaled so that the membrane voltages ranged between *E*_*L*_ = 0 and *ϑ* = 1. The scaling allowed weights of order 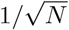, mitigating vanishing or exploding gradients during optimization. All optimization parameters are listed in Table 2 (Supplementary Materials).

**Table 2:** Parameter values related to optimization

| Symbol | Value / Init. Distribution | Dimensions | Description |
| --- | --- | --- | --- |
| $U$ | $\mathcal{U}(0.1, .25)$ | $N_E \times N_I$ | Initial release prob. |
| $W^{E \rightarrow I}$ | $\mathcal{N}(0, 1/N_E)$ | $N_E \times N_I$ | Exc. to Inh. weight |
| $W^{I \rightarrow I}$ | $\mathcal{N}(0, 1/N_I)$ | $N_I \times N_I$ | Inh. to Inh. weight |
| $W^{I \rightarrow D}$ | $\mathcal{N}(0, 0.2/N_I)$ | $N_I \times 1$ | Inh. to Exc. Dend. weight |
| $W^{I \rightarrow S}$ | $\mathcal{N}(0, 0.2/N_I)$ | $N_I \times 1$ | Inh. to Exc. Soma weight |
| - | 1e-3 | - | learning rate for weights |
| - | 4e-3 | - | learning rate for $U$ |
| $\beta$ | 10 | - | Slope spiking derivative |
| - | 1.0 | - | Gradient (absolute value) clipping |

### Optimization

We used gradient descent to find weights *W* and initial release probabilities *U* that minimize the difference between excitation and inhibition in both compartments:

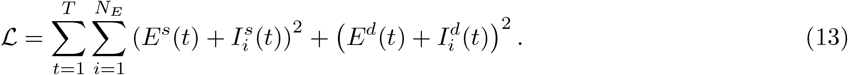

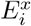 and 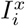 are the total excitatory and inhibitory input to compartment *x* ∈ {*s, d*} of PC *i*. To speed up the optimization process, all output synapses from a given neuron to a given compartment type had the same strength, i.e., the optimization of the output synapses is performed for *N*_*I*_ × 2 parameters. For the input synapses onto the INs, weight and initial release probability were optimized independent for all *N*_*E*_ × *N*_*I*_ synapses.

To achieve small interneuron rates necessary for interneuron specialization (Fig. A2), we subtracted the mean background input from 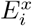:

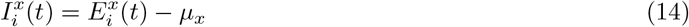

To propagate gradients through the spiking non-linearity, we replaced its derivative with the derivative of a smooth approximation [34]

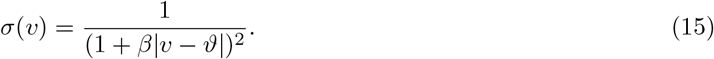

We used the machine learning framework PyTorch [68] to simulate the differential equations (forward Euler with step size 1 ms), compute the gradients of the objective ℒ using automatic differentiation, and update the network parameters using Adam [69]. The optimized parameters were initialized according to the distributions listed in Table 2 (Supplementary Materials). 2. We simulated the network response to batches of 8 trials of 600 ms, consisting of 100 ms pulses given at 2.5 Hz. The pulse amplitudes were drawn uniformly and independently for soma and dendrites from the set {100, 200, 300, 400}. Training converged within 200 batches (parameter updates). Before each parameter update, the gradient values were clipped between −1 and 1 to mitigate exploding gradients [70]. After each update, the initial release probability was clipped between 0 and 1 to avoid unphysiological values.

## Methods for Figures

### Figure 1

We measured the short-term plasticity of PC → IN synapses by simulating their response to two EPSPs given 10 ms apart, a typical interspike interval within a burst. The PPR was computed as the ratio of the two EPSP amplitudes, such that a PPR *>* 1 indicates short-term facilitation and a PPR *<* 1 indicates short-term depression. The PPR of a single IN was defined as the mean PPR of all its excitatory afferents. Clustering of interneurons was done by fitting a single Gaussian (before optimization) or a mixture of two Gaussians (after optimization) to the three-dimensional distribution of inhibitory weights to the PC soma, to PC dendrites, and the PC→IN Paired Pulse Ratio (PPR). Both models were fitted using Scikit-learn [71] on pooled data from five networks, trained from different random initializations. The density models where fitted on 246 interneurons that were active (firing rate higher than 1 spk/s) and had a medium to strong projection to either soma or dendrites (weight bigger than 0.01). The dashed lines in Fig. 1b illustrate the two-dimensional marginal distributions of the somatic and dendritic inhibition. All PCs received the same time-varying input currents, consisting of 100 ms pulses of 300 pA, given at a rate of 2.5 Hz. Correlations between compartment-specific excitation and inhibition were computed between the the currents to the PC compartments, averaged across all PCs in the network.

### Figure 2

Before optimization, we assigned interneurons to inhibit either PC somata or dendrites by fixing their weights onto the other compartment to zero. Half of the interneurons was assigned to inhibit the soma, the other half was assigned to inhibit the dendrites. Otherwise, weights and initial release probabilities were optimized as before.

### Figure 3

The definitions of burst rate, burst probability and event rate were taken from Naud & Sprekeler [33]: A burst was defined as multiple spikes occurring within 16 ms. The time of the first spike was taken as the time of the burst. An event was defined as a burst or a single spike. The instantaneous burst rate and event rate were computed by counting the number of bursts and events, respectively, in bins of 1ms and among the population of PCs, and smoothing the result with a Gaussian filter (width: 2ms). The burst probability was defined as

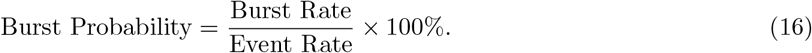

We injected current pulses of 100 ms duration to either soma or dendrite while injecting a constant current to the other compartment. Currents where varied in amplitude between 100 and 400 pA; the constant current was 0 pA. The figure shows the mean and standard deviation of the total network activity during 10 current pulses. For Fig. 3e, we injected simultaneous pulses to the other compartment of amplitude 0, 200 or 400 pA.

### Figure 4

We varied the correlation between the inputs to soma and dendrites by generating repeating current pulses with different temporal offsets and optimized a network for each offset. The interneuron specialization was defined as

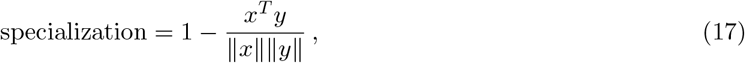

where *x* and *y* are *N*_*I*_ -dimensional vectors containing the inhibitory weights onto soma and dendrites and ‖ ·‖ the L_2_ norm. If each neuron inhibits either somata or dendrites, but not both, the specialization will be 1. If the weights are perfectly aligned (i.e., interneurons with a strong dendritic projection also have a strong somatic projection), the specialization will be 0. Here and in all figures, the EI correlation was computed as the correlation between the time series of the compartment-specific excitation and inhibition, after averaging across all PCs. Shown is the mean over 5 batches of 600 ms, where each batch consisted of 8 trials with amplitudes from {100, 200, 300, 400} pA, sampled independently for soma and dendrites.

### Figure 5

Figure 5a shows the connectivity strength over five networks. We first used the Gaussian mixture models to assign INs to PV or SST clusters, and then computed the mean connectivity between and within clusters for each network. For 5b, we trained networks with predefined interneuron populations to control the interneuron connectivity. Connections between populations were knocked out by fixing them to zero during and after optimization. EI correlations are computed for 5 batches of 600 ms, where each batch consisted of 8 trials with amplitudes from {100, 200, 300, 400} pA, sampled independently for soma and dendrites.

### Figure A.2

The minimum rate of PV neurons was controlled indirectly, by varying the baseline inhibitory target current to the soma—A larger baseline requires a higher minimum PV rate. We varied the minimum inhibitory current by subtracting only a fraction *α* of the baseline excitatory current:

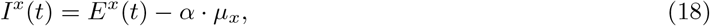

cf. Eq. (14). In the simulations, we varied *α* between 1 and 0.8, leading to a minimum PV rate between 1 spk/s, and 9 spk/s.

## A Supplementary Materials

### Supplementary Figures

**Figure A.1:**
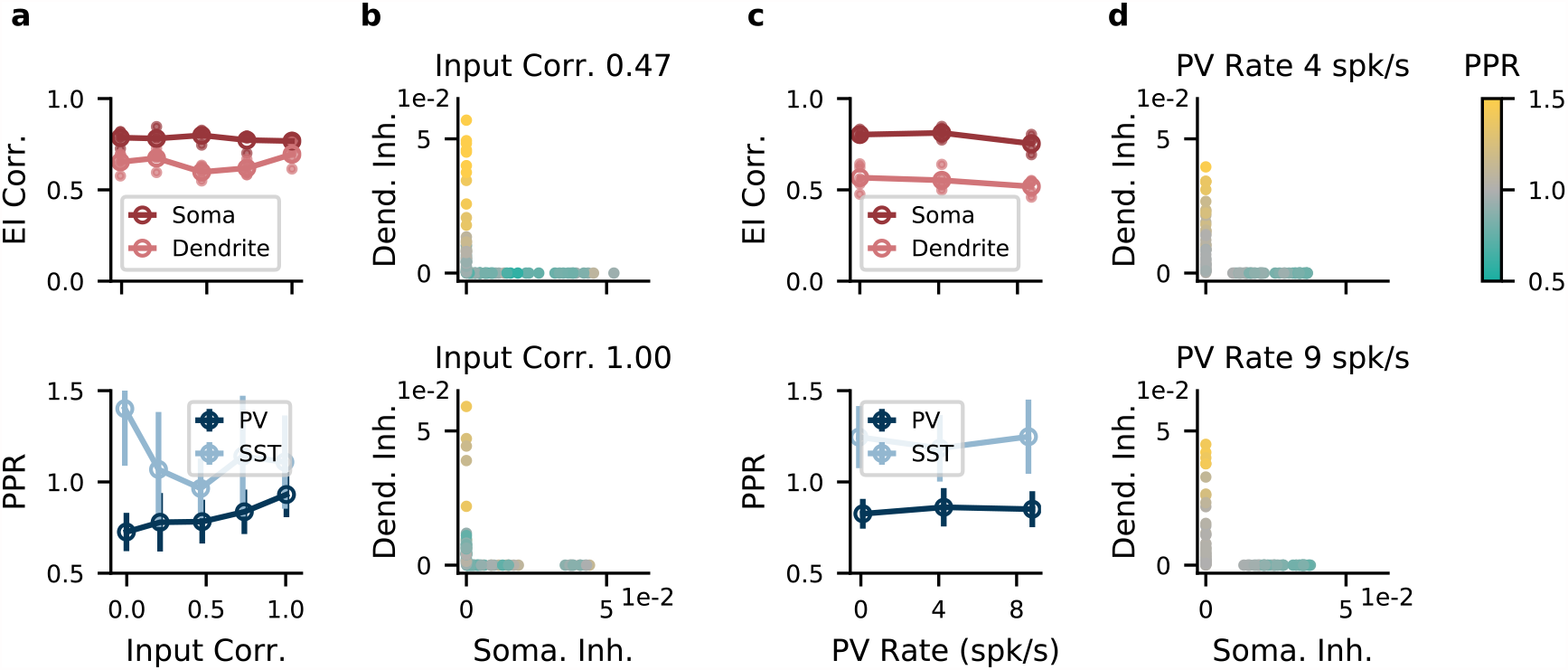
Non-overlapping interneuron populations achieve compartment-specific inhibition for a range of input statistics. **(a)** Top, performance as measured by compartment-specific correlation between excitation and inhibition of networks trained on different correlations between compartment-specific excitatory inputs. Open circles, mean over 5 batches of 8 stimuli with random amplitudes (see Methods). Small filled circles, individual batches. Here and in the other panels, the interneurons were assigned to inhibit only the soma or only the dendrites. Bottom, interneuron specialization as measured by Paired Pulse Ratio (PPR) decreases with input correlations. Error bars denote sd over IN populations. **(b)** Strength of somatic and dendritic inhibition from individual INs. Top, medium input correlation (0.47); bottom, high input correlation (1.00). Color indicates PPR. **c)** Top, as **a** but as function of minimum PV rate. Bottom, interneuron specialization as measured by Paired Pulse Ratio (PPR) is not influenced by minimum PV rate. Strength of somatic and dendritic inhibition from individual INs. Top, medium PV rate (4 spk/s); bottom, high PV rate (9 spk/s).

**Figure A.2:**
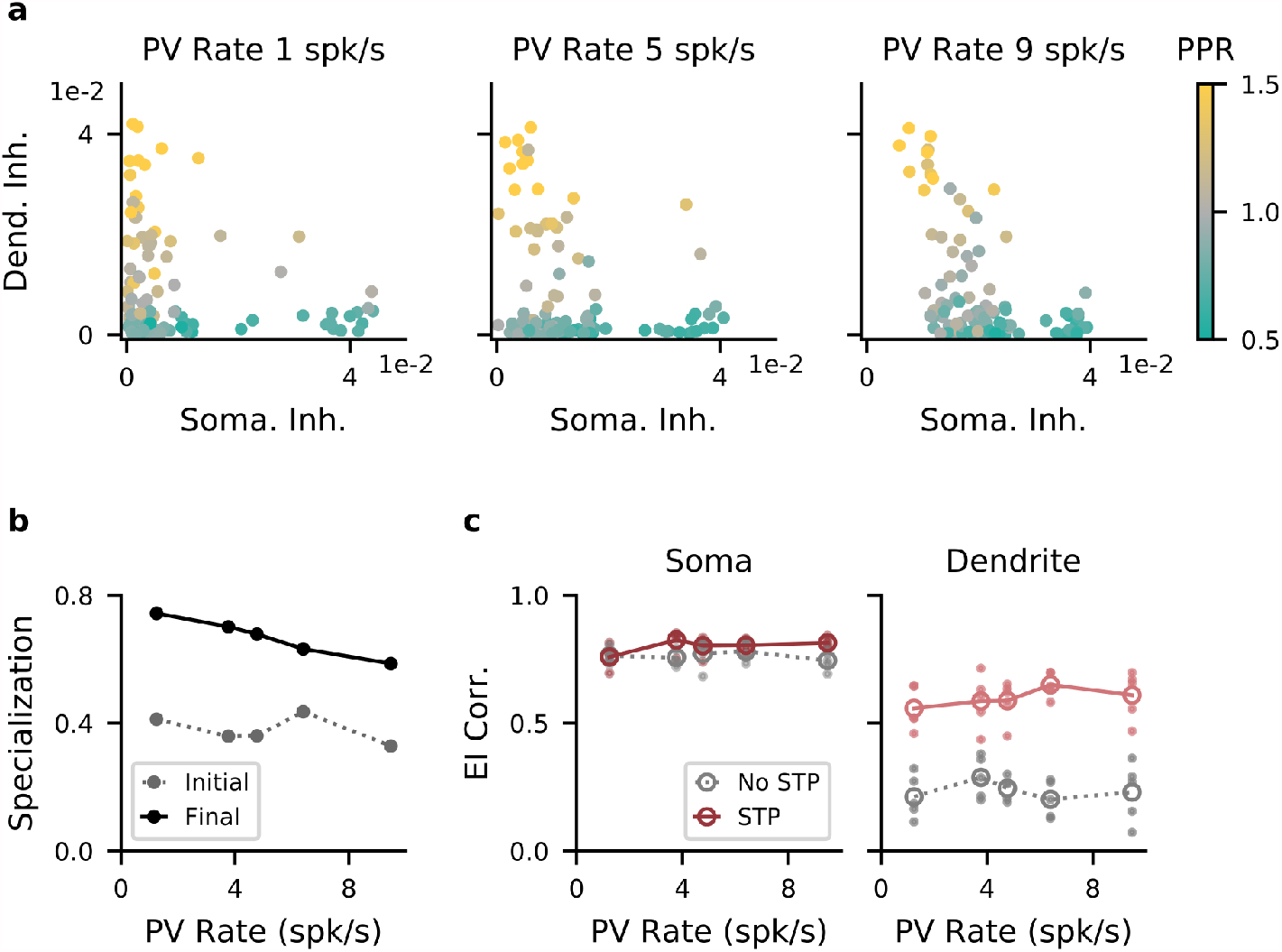
Higher baseline PV rates decrease the need for interneuron specialization. **(a)** Strength of somatic and dendritic inhibition from individual INs. Left, middle, right: network optimized with a baseline PV rate of 1 (low), 5 (medium), and 9 spk/s (high), respectively. **(b)** Specialization of IN → E weights. If each IN targets either soma or dendrites, the specialization is 1 (see Methods). Gray: specialization of initial, random network; black: specialization after optimization. **(c)** Left, correlation between excitation and inhibition as function of minimum PV rate. Red: networks with optimized short-term plasticity. Gray: Networks without short-term plasticity. Open circles, mean over 5 batches of 8 stimuli with random amplitudes. Small filled circles, individual batches.

### Mathematical Analysis of a Simplified Network Model

We performed a mathematical analysis of a simplified network to better understand the following results of our spiking network simulations:

1. A compartment-specific balance requires PV → SST inhibition, but no other IN → IN connectivity (Fig. 5).
2. Higher interneuron rates require less IN specialization, i.e., individual interneurons often inhibit both PC compartments (Fig A.2).

The simplified model consists of a population of principal cells (PC) and two populations of interneurons that we will refer to as parvalbumin (PV)-positive and as somatostatin (SST)-positive cells. The population activity of the PCs is represented by somatic activity *e* and dendritic activity *b*. The interneuron activities are represented by firing rates *p* and *s*. The four activity variables *e, b, p, s* are best thought of as deviations of the respective activity from baseline. The activity variables can hence be both positive and negative (ignoring saturation effects that arise when the baseline is very low, see below).

For our analysis, we make the following assumptions: (1) somatic input linearly increases somatic activity *e*, (2) dendritic input linearly increases dendritic activity *b*, which is in turn assumed to be independent of somatic input/activity (note that the latter assumption deviates from a BAC-firing mechanism [31], but is necessary to obtain a linear model), (3) the activities *p, s* of the interneuron populations increase linearly with their input, and (4) short-term plasticity is characterized by a single, static parameter (see below). Because we are interested only in qualitative statements, the analysis is done in terms of unitless variables. The model describes the dynamics of the four activity variables *e, b, p, s*:

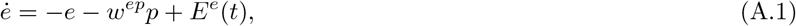

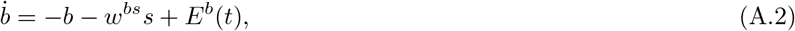

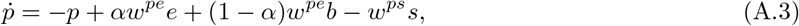

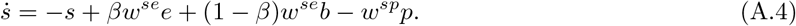

Here, the synaptic weight from population *y* to *x* is modeled with a non-negative weight *w*^*xy*^ (*x, y* ∈ {*e, p, s*}; *e*: PCs, *p*: PV INs, *s*: SST INs). The central tenet of this simplified model is that somatic and dendritic activity both generate characteristic spike patterns in PCs—such as events and bursts—which are selectively transmitted by synapses because of short-term plasticity. The parameters *α, β* ∈ [0, 1] describe the short-term plasticity of the PC→PV and PC→SST synapses, respectively. *α, β* = 1 corresponds to synapses that only transmit somatic activity. If somatic activity generates events and dendritic activity generates bursts, this would require “perfectly depressing” synapses, i.e., synapses that transmit only the first spike of a burst. *α, β* = 0 corresponds to synapses that only transmit dendritic activity. For the case where dendritic activity generates bursts, this requires “perfectly facilitating” synapses that ignore individual spikes and transmit only bursts. We assumed that the projections of the interneurons are specialized, i.e., that PV interneurons inhibit the soma and SST interneurons inhibit the dendrite. We will abandon this assumption in Section A.1. We also excluded inhibitory recurrence within the two populations (PV → PV, SST → SST), because these connections would only change the effective time constant of the respective activation variable. The somata and dendrites of the PCs receive time-varying external inputs *E*^*e*^(*t*) and *E*^*b*^(*t*), respectively. All activity variables follow leaky dynamics.

The dynamical system can be written as 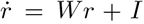, where the vector *r* contains the activation variables *r* = (*e, b, p, s*)^*T*^, *I* contains the external inputs *I* = (*E*^*e*^, *E*^*b*^, 0, 0)^*T*^, and *W* is the matrix of effective connectivity strengths

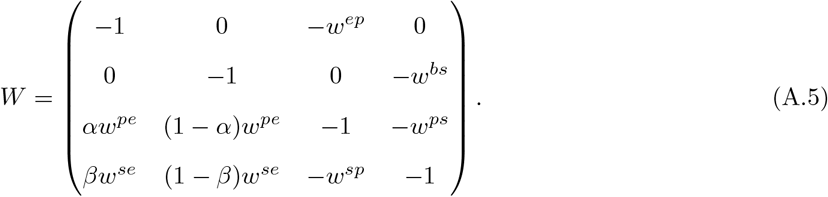

Assuming that the time constant of the network is sufficiently short to adiabatically follow the input currents, we can consider the steady state by setting 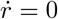 and solving for *r*:

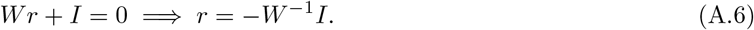

### Influence of IN→IN connections on compartment-specific E/I balance

In the steady state Eq. (A.6), the IN rates are equal to

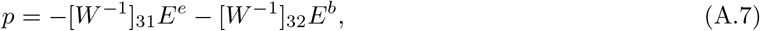

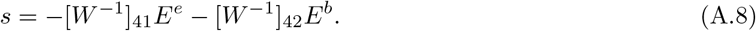

Here, [*W*^*−*1^]_*ij*_ refers to the element in row *i* and column *j* of the matrix *W*^*−*1^. Assuming that the interneurons specialize by inhibiting a single compartment, a necessary (and, up to scaling, sufficient) condition for compartment-specific balance is that the PV rate *p* is proportional to the external input targeting the soma and independent of the input targeting the dendrite. Similarly, the SST rate should be proportional to the external input targeting the dendrite and independent of the input targeting the soma. By Eq. (A.7), a compartment-specific balance hence requires [*W*^*−*1^]_32_ = 0 and [*W*^*−*1^]_41_ = 0. Computing these matrix entries yields:

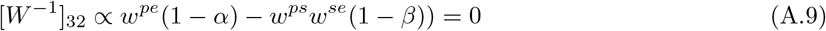

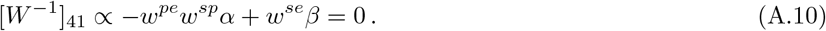

These equations have a simple interpretation. Each of the two terms in [*W*^*−*1^]_32_ represents a pathway by which dendritic activity reaches the PV interneurons. The first term quantifies how much dendritic activity reaches PV interneurons via the direct excitatory PC→PV projection, the second represents corresponding feedforward inhibition via the PC→SST→PV pathway. If these two pathways cancel, PV activity is independent of dendritic activity. Similarly, the two pathways in [*W*^*−*1^]_41_ that transmit somatic activity to the SST need to cancel.

What is the role of short-term plasticity? For illustration, let us first consider the limiting case of “perfect” synaptic depression (*α* = 1). Perfectly depressing PC → PV synapses would imply that the PV interneurons only receive somatic activity from the PCs via the direct PC→PV pathway. The condition (A.9) then reduces to

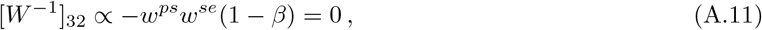

i.e., dendritic activity should not reach PV interneurons via the indirect PC→SST→PV pathway, because this would render PV activity dependent on dendritic activity. Because dendritic activity need to be transmitted to the SST interneurons to reach an E/I balance in the dendrite, this implies that the SST→PV connection should be absent.

“Perfect” synaptic depression (*α, β* = 1) or facilitation (*α, β* = 0) are hard to implement, certainly by a Markram-Tsodyks model in the presence of background activity. However, the effect of imperfect depression in the PC→PV connection (*α <* 1) can be compensated by feedforward inhibition along the PC→SST→PV pathway. Similarly, imperfect PC→SST facilitation picks up somatic activity, which can then be canceled by feedforward inhibition via the PC→PV→SST pathway. The role of IN→IN synapses is therefore to complement “imperfect” short-term plasticity in decoding compartment-specific inputs.

The observation that PV→SST connections are the most important IN→IN connections in our model results from “imperfect” facilitation in the excitatory synapses onto SST interneurons. Because events occur more frequently than bursts, the excess excitation they trigger in SST interneurons needs to be actively cancelled via the PV→SST pathway. The converse SST→PV connection is less critical, because bursts are comparatively rare, such that their transmission via PC→PV synapses causes only minor disturbances of the compartment-specific E/I balance.

### A.1 Influence of IN baseline firing rates on interneuron specialization

The previous analysis assumed that interneurons were specialized to inhibit a single compartment. When should we expect specialization in the first place? We can investigate this question by extending the simplified model by inhibition from all INs onto all PC compartments:

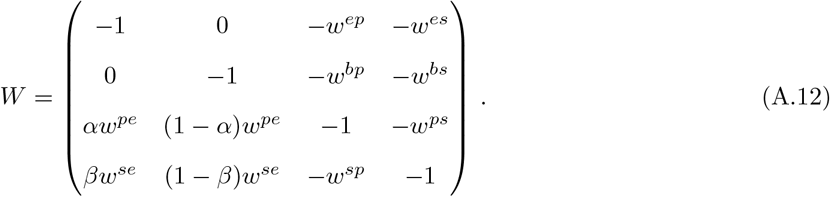

A compartment-specific balance now requires external input to be canceled by the inhibition from both interneurons:

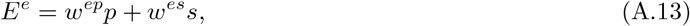

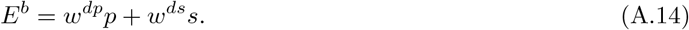

Without additional constraints, this system has an infinite number of solutions, i.e., weight configurations that achieve a compartment-specific balance. However, the simple constraint of low baseline firing rates of the interneurons collapses the solution space to the specialized one (*w*^*es*^ = *w*^*dp*^ = 0), for the following reason.

The activity variables *p, s* represent deviations of the interneurons firing rates from baseline. If the baseline is sufficiently high, these deviations can be both positive and negative. In that case, inhibition from one interneuron class can be cancelled by disinhibition from the other interneuron class. PV and SST interneurons are then both free to respond to both somatic and dendritic activity, as long as the weighted sums of the inhibition and disinhibition they provide to PC somata and dendrites mirrors the excitatory input to those compartments. There are many ways of doing so.

For low baseline firing rates, disinhibition is no longer available, because negative deviations from baseline are limited by the fact that activities cannot be negative. For illustration, let us consider the case where both PV and SST neurons have zero baseline activity. By definition, the excitatory signals *E*^*e/b*^ are zero for baseline activity, because they represent deviations from baseline. Therefore, we can assume that there exists a moment where the input to one PC compartment is zero, while the input to the other compartment is positive (e.g., *E*^*e*^ = 0, *E*^*b*^ *>* 0). By Eq. (A.13) and because all weights and rates must be positive, *w*^*ep*^*p* + *w*^*es*^*s* = 0 implies that *w*^*ep*^ = 0 or *p* = 0 and *w*^*es*^ = 0 or *s* = 0. At least one weight has to be non-zero (otherwise balancing the soma is impossible), and at least one rate has to be non-zero (otherwise balancing the dendrite is impossible). Without loss of generality we can conclude that *w*^*ep*^ *>* 0, *p* = 0, *w*^*es*^ = 0, and *s >* 0: The PC soma is only inhibited by the PV neuron. Analogously, the existence of a moment when *E*^*b*^ = 0 but *E*^*e*^ *>* 0 implies that *w*^*ds*^ *>* 0, *w*^*dp*^ = 0, meaning that the PC dendrite is only inhibited by the SST neuron. If the baseline activity is low, but not strictly zero, this saturation arguments still hold, if the variations in the firing rates that are required to balance the external input are larger than the baseline activity. Low baseline firing rates therefore imply interneuron specialization, because they prevent inhibition and disinhibition from non-specialized neurons to cancel.

